# A deep learning framework for predicting the neutralizing activity of COVID-19 therapeutics and vaccines against evolving SARS-CoV-2 variants

**DOI:** 10.1101/2023.10.24.563847

**Authors:** Robert P. Matson, Isin Y. Comba, Eli Silvert, Michiel J.M. Niesen, Karthik Murugadoss, Dhruti Padwardhan, Rohit Suratekar, Elizabeth-Grace Goel, Brittany J. Poelaert, Kanny Wan, Kyle R. Brimacombe, AJ Venkatakrishnan, Venky Soundararajan

## Abstract

Understanding how viral variants evade neutralization is crucial for improving antibody-based treatments, especially with rapidly evolving viruses like SARS-CoV-2. Yet, conventional assays are limited in the face of rapid viral evolution, relying on a narrow set of viral isolates, and falling short in capturing the full spectrum of variants. To address this, we have developed a deep learning approach to predict changes in neutralizing antibody activity of COVID-19 therapeutics and vaccines against emerging viral variants. First, we trained a variational autoencoder (VAE) using all 67,885 unique SARS-CoV-2 spike protein sequences from the NCBI virus (up to October 31, 2022) database to encode spike protein variants into a latent space. Using this VAE and a curated dataset of 7,069 *in vitro* assay data points from the NCATS OpenData Portal, we trained a neural network regression model to predict fold changes in neutralizing activity of 40 COVID-19 therapeutics and vaccines against spike protein sequence variants, relative to their neutralizing activity against the ancestral strain (Wuhan-Hu-1). Our model also employs Bayesian inference to quantify prediction uncertainty, providing more nuanced and informative estimates. To validate the model’s predictive capacity, we assessed its performance on a test set of *in vitro* assay data collected up to eight months after the data included in the model training (N = 980). The model accurately predicted fold changes in neutralizing activity for this prospective dataset, with an R^2^ of 0.77. Expanding our methodology to include all available data from NCBI virus and NCATS OpenData Portal up to date, we assessed predicted changes in activity for current COVID-19 monoclonal antibodies and vaccines against newly identified SARS-CoV-2 lineages. Our predictions suggest that current therapeutic and vaccine-induced antibodies will have significantly reduced activity against newer XBB descendants, notably EG.5, FL.1.5.1, and XBB.1.16. Using the model, we were able to primarily attribute the observed predicted loss in activity to the F456L spike mutation found in EG.5 and FL.1.5.1 sequences. Conversely, mRNA-bivalent vaccines are predicted to be less susceptible to the recent BA.2.86 variant compared to new XBB descendants. These findings align closely with recent research, underscoring the potential of deep learning in shaping therapeutic and vaccine strategies for emerging viral variants.

## Introduction

Viruses can accumulate sequence changes under immune selection pressure and due to natural genetic variation.^1–3^ Such mutations can permit evasion of host immune responses, leading to the emergence of new viral variants that reduce the efficacy of vaccines and antibody-based treatments.^4,5^ With the ongoing evolution of a virus, there arises an uncertainty as to whether antibodies induced by therapies and vaccines will be effective in identifying and neutralizing novel strains. Because of this, it is crucial to monitor viral strains’ potential for antibody escape to revise clinical and public health guidelines and develop more effective therapeutic and vaccine strategies.

Cell-based assays are a widely used tool for assessing the antibody evasion potential of viral strains.^6,7^ These assays involve exposing a viral strain to an antibody agent in cell culture and evaluating the level of viral replication, infectivity, or virulence *in vivo* or *in vitro*. However, these assays have certain limitations, particularly when the virus is evolving rapidly. This is due to the fact that they rely on a limited number of viral isolates, which may not adequately represent the full diversity of circulating strains.^8–10^ Consequently, it becomes challenging to monitor viral escape entirely and develop effective treatments that can target a wide range of viral strains. This challenge has been particularly evident during the COVID-19 pandemic, as the SARS-CoV-2 virus continues to evolve and produce new lineages and sub-lineages, with over 1.7 million unique sequences recorded to date.^11^ Therefore, it is essential to complement these assays with surveillance efforts and realistic models to ensure that emerging viral strains and underlying antibody escape properties are entirely detected and monitored in real-time.

Deep learning, which is a subfield of machine learning, has the potential to provide a complementary approach in monitoring the antibody evasion potential of emerging viral variants. Due to its ability to rapidly extrapolate information from complex biological data, deep learning has emerged as a favorable method in the biomedical field.^12–14^ Recent studies have successfully modeled the temporal and geographic evolution of SARS-CoV-2 and predicted the impact of mutations on ACE2 binding using deep learning methods.^15–17^ The arrival of new influenza strains each year has also driven the development of deep learning models that are useful in the creation of annual flu vaccines.^18–20^ Furthermore, in an era where there is a significant amount of viral surveillance data,^21–23^ deep learning could be used to extract valuable knowledge and insights about antibody evasion that may not be entirely captured by conventional assays.

Here, we propose a deep learning method to predict changes in neutralizing antibody activity of COVID-19 therapeutics and vaccines against emerging SARS-CoV-2 variants. Our method utilizes a variational autoencoder (VAE) to encode spike protein sequences into a latent space embedding, allowing viral sequences to be inputted into a predictive model. Using compiled *in vitro* assay data, we trained a neural network model to predict fold changes in the neutralization activity of COVID-19 therapeutics and vaccines against spike protein variants, relative to their activity against the ancestral strain (Wuhan-Hu-1). This work presents a comprehensive analysis of the spike protein variants and corresponding antibody resistance of SARS-CoV-2, augmenting the insights derived from experimental assays. Through this research, advancements can be made towards developing more effective therapeutic and vaccine strategies against rapidly evolving viruses. Additionally, it can facilitate the detection of viral variants that may evade current approved treatments and the discovery of antibodies that have regained their effectiveness against new variants.

## Materials and methods

### Variational autoencoder (VAE) architecture, training, and optimization

SARS-CoV-2 genomes present in the NCBI Virus Database were downloaded. Sequences collected up to October 31, 2022, were used to train the VAE. The dataset consisted of 1,208,321 spike protein amino acid sequences. We aligned and translated the sequences using NextClade.^24^ Duplicate spike protein sequences were removed, which resulted in a final dataset of 67,885 unique spike protein sequences. The sequences were one-hot encoded as 22 by 1,273 arrays, as there are 1,273 amino acid positions in the Wuhan-Hu-1 spike protein and 22 options at each site: any one of the 20 amino acids, an insertion, or a deletion. In the case of an insertion, we only encoded that there was an insertion following the current position, without encoding the identity of the inserted fragment.

The VAE consists of an encoder and a decoder. The encoder compresses the spike protein sequence data (one-hot encoded amino acid sequences) to its latent embedding and the decoder reconstructs the input sequence data from its latent embedding. In the encoder, the number of latent dimensions is set to 32 because additional gains in loss after decoding were minimal when increasing the number of latent dimensions beyond 32. The latent space is modeled as a multivariate normal distribution with a defined latent mean and variance (log-transformed for numerical stability). A standard normal distribution was used as the prior distribution for the latent space. A sampling layer is present in the encoder, where data is randomly sampled from the latent space distribution before being passed to the decoder.

The VAE was compiled with an Adam optimizer. The loss function for the VAE is the sum of the reconstruction loss and the Kullback-Leibler (KL) divergence between the distribution returned by the encoder and a standard normal distribution. The reconstruction loss is calculated using binary cross-entropy. The VAE loss function is defined as follows:

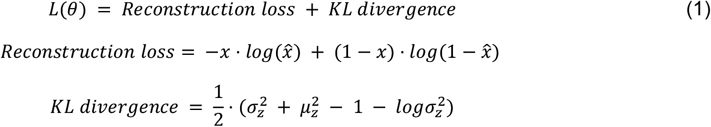

where *x* represents the input data (one-hot encoded amino acid sequences), 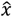 represents the reconstructed data, 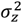 represents the variance of the latent distribution, and *μ*_*z*_ represents the mean of the latent distribution. The reconstruction loss quantifies the ability to reconstruct the input sequence data from its latent embedding, while the KL-divergence ensures that the learned latent space distribution follows a standard normal distribution. This helps the VAE learn smooth latent representations where distance in latent space reflects similarity between inputted sequences. The VAE was trained on 80% of the sampled sequences (N = 54,308) and evaluated on the remaining 20% of sampled sequences (N = 13,577). We trained the VAE for 50 epochs with a batch size of 32, while also providing an early stopping function with a patience of five to stop training once the validation loss stopped improving for five consecutive epochs.

### OpenData Portal curation of neutralizing activity against SARS-CoV-2 variants

The SARS-CoV-2 variant therapeutic data on the OpenData Portal have been curated by the National Center for Advancing Translational Sciences (NCATS) in collaboration with the National Institutes of Health (NIH) Accelerating COVID-19 Therapeutic Interventions and Vaccines (ACTIV) Preclinical Working Group and Tracking Resistance and Coronavirus Evolution (TRACE) initiative with support from the Foundation for the National Institutes of Health (FNIH). The data shared on the OpenData Portal have been manually curated from publications (both preprints and peer-reviewed articles) or data directly submitted. Curation efforts prioritized publications on advanced stage therapeutic agents (those approved and/or in clinical trials), with an emphasis on studies conducted 1) by each agent’s parent pharmaceutical company or 2) with a government partner. Curation efforts collated neutralizing activity of vaccines and therapeutics against SARS-CoV-2 variants, in addition to related metadata, from *in vitro* assays utilizing live or pseudotyped viruses. Fold-reductions and raw values provided via publication or direct submission can be accessed via download on the OpenData Portal web browser (https://opendata.ncats.nih.gov/variant/activity).

### Preprocessing and encoding of in vitro neutralization assay data

SARS-CoV-2 *in vitro* neutralization activity data compiled by the National Center for Advancing Translational Sciences (NCATS) on its COVID-19 OpenData Portal^25^ were used for model development. The dataset consists of *in vitro* assays collected between January 9, 2021, and June 22, 2023. Both live virus replication assays and pseudotyped virus assays were considered for this analysis. Neutralization activity fold change ratios between the SARS-CoV-2 wild-type strain (Wuhan-Hu-1) and spike protein variants were compiled and log10 transformed. COVID-19 therapeutics and vaccines with less than 20 data points were excluded from our study to ensure high confidence in the predictions. Spike protein sequences from variants tested were auto-encoded into 32 latent dimensions using the previously described VAE. The specific therapeutics and vaccines evaluated in each assay were one-hot encoded into 40 additional dimensions, consisting of 19 monoclonal antibodies, 11 vaccines (vaccine sera samples) and 10 convalescent plasma samples (**Table 1**). To enable quantification of performance against future data, we divided the available data into training and test sets based on the data collection date. Specifically, the training set consisted of sequences tested between January 9, 2021, and October 31, 2022 (N = 6,089), and the remaining test set consisted of sequences tested between November 1, 2022, and June 22, 2023 (N = 980). **Table 2** reports the number and percentage of sequences belonging to a particular SARS-CoV-2 viral lineage in both the training and test sets.

**Table 1:** Number (percentage) of COVID-19 therapeutic agents in neural network model training and test sets. Rows are colored by the therapeutic class.

| Therapeutic class | Therapeutic agent | Training set | Test set | All |
| --- | --- | --- | --- | --- |
| Convalescent plasma | B.1.1.529 Conv. Plasma/Sera | 55 (0.9%) | 0 (0.0%) | 55 (0.8%) |
| Convalescent plasma | B.1.1.7 Conv. Plasma/Sera | 30 (0.5%) | 0 (0.0%) | 30 (0.4%) |
| Convalescent plasma | B.1.351 Conv. Plasma/Sera | 31 (0.5%) | 0 (0.0%) | 31 (0.4%) |
| Convalescent plasma | B.1.617.2 Conv. Plasma/Sera | 52 (0.9%) | 13 (1.3%) | 65 (0.9%) |
| Convalescent plasma | BA.1 Conv. Plasma/Sera | 63 (1.0%) | 7 (0.7%) | 70 (1.0%) |
| Convalescent plasma | BA.2 Conv. Plasma/Sera | 20 (0.3%) | 54 (5.5%) | 74 (1.0%) |
| Convalescent plasma | BA.4/5 Conv. Plasma/Sera | 4 (0.1%) | 25 (2.6%) | 29 (0.4%) |
| Convalescent plasma | BA.5 Conv. Plasma/Sera | 0 (0.0%) | 26 (2.7%) | 26 (0.4%) |
| Convalescent plasma | COVID-19 Conv. Plasma/Sera | 179 (2.9%) | 0 (0.0%) | 179 (2.5%) |
| Convalescent plasma | COVID-HIG | 39 (0.6%) | 0 (0.0%) | 39 (0.6%) |
| Monoclonal antibody | Adintrevimab | 69 (1.1%) | 0 (0.0%) | 69 (1.0%) |
| Monoclonal antibody | Amubarvimab | 165 (2.7%) | 31 (3.2%) | 196 (2.8%) |
| Monoclonal antibody | Amubarvimab+Romlusevimab | 28 (0.5%) | 18 (1.8%) | 46 (0.7%) |
| Monoclonal antibody | Bamlanivimab | 233 (3.8%) | 0 (0.0%) | 233 (3.3%) |
| Monoclonal antibody | Bamlanivimab+Etesevimab | 61 (1.0%) | 0 (0.0%) | 61 (0.9%) |
| Monoclonal antibody | Bebtelovimab | 402 (6.6%) | 58 (5.9%) | 460 (6.5%) |
| Monoclonal antibody | Casirivimab | 369 (6.1%) | 3 (0.3%) | 372 (5.3%) |
| Monoclonal antibody | Cilgavimab | 580 (9.5%) | 56 (5.7%) | 636 (9.0%) |
| Monoclonal antibody | Etesevimab | 233 (3.8%) | 0 (0.0%) | 233 (3.3%) |
| Monoclonal antibody | Evusheld | 482 (7.9%) | 56 (5.7%) | 538 (7.6%) |
| Monoclonal antibody | Imdevimab | 332 (5.5%) | 3 (0.3%) | 335 (4.7%) |
| Monoclonal antibody | Regdanvimab | 70 (1.1%) | 0 (0.0%) | 70 (1.0%) |
| Monoclonal antibody | Romlusevimab | 119 (2.0%) | 18 (1.8%) | 137 (1.9%) |
| Monoclonal antibody | Ronapreve | 78 (1.3%) | 3 (0.3%) | 81 (1.1%) |
| Monoclonal antibody | S2E12 | 38 (0.6%) | 18 (1.8%) | 56 (0.8%) |
| Monoclonal antibody | S309 | 170 (2.8%) | 61 (6.2%) | 231 (3.3%) |
| Monoclonal antibody | Sotrovimab | 344 (5.6%) | 5 (0.5%) | 349 (4.9%) |
| Monoclonal antibody | Tixagevimab | 610 (10.0%) | 58 (5.9%) | 668 (9.4%) |
| Monoclonal antibody | VIR-7832 | 26 (0.4%) | 0 (0.0%) | 26 (0.4%) |
| Vaccine | Ad26.COV2.S | 73 (1.2%) | 0 (0.0%) | 73 (1.0%) |
| Vaccine | BNT162b2 | 233 (3.8%) | 34 (3.5%) | 267 (3.8%) |
| Vaccine | Bivalent BNT162b2 OMI (BA.4/5) | 4 (0.1%) | 21 (2.1%) | 25 (0.4%) |
| Vaccine | ChAdOx1 nCoV-19 | 35 (0.6%) | 2 (0.2%) | 37 (0.5%) |
| Vaccine | CoronaVac | 416 (6.8%) | 228 (23.3%) | 644 (9.1%) |
| Vaccine | Unspecified/Mixed Bivalent Vaccine | 0 (0.0%) | 72 (7.3%) | 72 (1.0%) |
| Vaccine | Unspecified/Mixed Vaccine [Prime + Boost] | 132 (2.2%) | 102 (10.4%) | 234 (3.3%) |
| Vaccine | Unspecified/Mixed Vaccine [Prime] | 26 (0.4%) | 0 (0.0%) | 26 (0.4%) |
| Vaccine | ZF2001 | 33 (0.5%) | 0 (0.0%) | 33 (0.5%) |
| Vaccine | mRNA-1273 | 231 (3.8%) | 8 (0.8%) | 239 (3.4%) |
| Vaccine | mRNA-1273.351 | 24 (0.4%) | 0 (0.0%) | 24 (0.3%) |

**Table 2:** Number (percentage) of SARS-CoV-2 lineages in neural network model training and test sets.

| Viral lineage | Training set | Test set | All |
| --- | --- | --- | --- |
| Single mutation | 2717 (44.6%) | 0 (0.0%) | 2717 (38.4%) |
| BA.2 | 469 (7.7%) | 534 (54.5%) | 1003 (14.2%) |
| B.1.351 | 553 (9.1%) | 2 (0.2%) | 555 (7.9%) |
| Omicron: Other | 407 (6.7%) | 41 (4.2%) | 448 (6.3%) |
| BA.4/5 | 178 (2.9%) | 241 (24.6%) | 419 (5.9%) |
| B.1.1.7 | 380 (6.2%) | 0 (0.0%) | 380 (5.4%) |
| B.1.617.2 | 316 (5.2%) | 2 (0.2%) | 318 (4.5%) |
| B.1.1.529 | 299 (4.9%) | 0 (0.0%) | 299 (4.2%) |
| P.1 | 216 (3.5%) | 0 (0.0%) | 216 (3.1%) |
| B.1.617.1 | 92 (1.5%) | 0 (0.0%) | 92 (1.3%) |
| B.1.427/429 | 88 (1.4%) | 0 (0.0%) | 88 (1.2%) |
| B.1.526 | 64 (1.1%) | 0 (0.0%) | 64 (0.9%) |
| BQ.1.1 | 8 (0.1%) | 54 (5.5%) | 62 (0.9%) |
| BQ.1 | 8 (0.1%) | 23 (2.3%) | 31 (0.4%) |
| B.1.429 | 30 (0.5%) | 0 (0.0%) | 30 (0.4%) |
| XBB | 4 (0.1%) | 25 (2.6%) | 29 (0.4%) |
| B.1.621 | 27 (0.4%) | 0 (0.0%) | 27 (0.4%) |
| C.37 | 27 (0.4%) | 0 (0.0%) | 27 (0.4%) |
| B.1.525 | 25 (0.4%) | 0 (0.0%) | 25 (0.4%) |
| BN.1 | 6 (0.1%) | 17 (1.7%) | 23 (0.3%) |
| CH.1.1 | 0 (0.0%) | 17 (1.7%) | 17 (0.2%) |
| B.1.1.519 | 16 (0.3%) | 0 (0.0%) | 16 (0.2%) |
| B.1.616 | 14 (0.2%) | 0 (0.0%) | 14 (0.2%) |
| AY | 14 (0.2%) | 0 (0.0%) | 14 (0.2%) |
| XBF | 0 (0.0%) | 12 (1.2%) | 12 (0.2%) |
| CJ.1.1 | 0 (0.0%) | 12 (1.2%) | 12 (0.2%) |
| P.2 | 11 (0.2%) | 0 (0.0%) | 11 (0.2%) |
| B.1.2 | 9 (0.1%) | 0 (0.0%) | 9 (0.1%) |
| B.1.1.298 | 9 (0.1%) | 0 (0.0%) | 9 (0.1%) |
| C.1.2 | 8 (0.1%) | 0 (0.0%) | 8 (0.1%) |
| A.VOI.V2 | 7 (0.1%) | 0 (0.0%) | 7 (0.1%) |
| R.1 | 6 (0.1%) | 0 (0.0%) | 6 (0.1%) |
| A.27 | 6 (0.1%) | 0 (0.0%) | 6 (0.1%) |
| A.23.1 | 6 (0.1%) | 0 (0.0%) | 6 (0.1%) |
| B.1.351, P.1 | 6 (0.1%) | 0 (0.0%) | 6 (0.1%) |
| B.1.1.33, B.1.1.1 | 6 (0.1%) | 0 (0.0%) | 6 (0.1%) |
| B | 6 (0.1%) | 0 (0.0%) | 6 (0.1%) |
| B.1.214.2 | 6 (0.1%) | 0 (0.0%) | 6 (0.1%) |
| A.23 | 5 (0.1%) | 0 (0.0%) | 5 (0.1%) |
| B.1.177.31 | 5 (0.1%) | 0 (0.0%) | 5 (0.1%) |
| B.1.388 | 4 (0.1%) | 0 (0.0%) | 4 (0.1%) |
| B.1.258 | 4 (0.1%) | 0 (0.0%) | 4 (0.1%) |
| AV.1 | 4 (0.1%) | 0 (0.0%) | 4 (0.1%) |
| R.2 | 4 (0.1%) | 0 (0.0%) | 4 (0.1%) |
| Multiple Mutations | 4 (0.1%) | 0 (0.0%) | 4 (0.1%) |
| B.1.523 | 3 (0.0%) | 0 (0.0%) | 3 (0.0%) |
| B.1.619.1 | 3 (0.0%) | 0 (0.0%) | 3 (0.0%) |
| B.1.617 | 3 (0.0%) | 0 (0.0%) | 3 (0.0%) |
| B.1.625 | 3 (0.0%) | 0 (0.0%) | 3 (0.0%) |
| C.36 | 2 (0.0%) | 0 (0.0%) | 2 (0.0%) |
| B.1.619 | 1 (0.0%) | 0 (0.0%) | 1 (0.0%) |

### Uncertainty-based neural network model architecture, training, and optimization

A neural network model underwent training, hyperparameter tuning, and cross-validation of the training set to predict fold changes in neutralization activity for COVID-19 therapeutics and vaccines against variant spike protein sequences. The model was compiled with an Adam optimizer and a custom loss function (Equation 2).^26^ The activation function for the input and hidden layers is a leaky rectified linear unit function (LReLU) and the activation function for the output layer is a linear function. We trained the model for 100 epochs with a batch size of 32, while also applying an early stopping function with a patience of ten to stop training once the validation loss stopped improving for ten consecutive epochs.

A custom loss function, based on prior literature, was used to optimize our neural network (Equation 2), where the output layer of the model consists of two neurons to predict the mean and variance of each observation.^26^ The loss function can be described as:

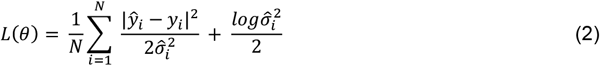

Where *ŷ*_*i*_ is the prediction for the *i*-th observation, 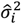 is the estimated variance for the *i*-th observation, and *y*_*i*_ is the true target value for the *i*-th observation. Typically, a Bayesian neural network is trained to predict the log variance, 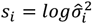 (Equation 3):

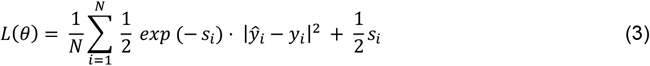

This is because the loss avoids a potential division by zero.^26^ The exponential mapping also allows us to regress unconstrained scalar values, where *exp*(−*s*_*i*_) is resolved to the positive domain giving valid values for variance.^26^ Non-Bayesian neural network training methods usually ignore the variance term in this equation, assuming it is constant across all observations in the data. However, by adding the variance term, the variance can be implicitly learned as a function of the data and can be used as a measure of uncertainty inherent in the observations. Including the variance term also allows the model to be more robust to noisy data because observations with higher variance (i.e., higher uncertainty) will have a smaller effect on the loss.

## Results

### Encoding SARS-CoV-2 spike protein sequences using VAE

A VAE was first developed to encode SARS-CoV-2 spike protein sequences and create a latent space that captures mutational patterns and relationships between sequences. The dataset comprised 67,885 unique spike protein sequences extracted from the NCBI Virus Database as of October 31, 2022 (**Figure 1a**). To train the VAE, 54,308 sequences were fed into the encoder, which compressed them into a 32-dimensional latent space (**Figure 1b, Figure 1c**). The decoder then reconstructed the sequences from their latent embedding. Following training, a difference score was calculated between the input and output sequences, indicating how well the decoder reconstructed the sequences from their latent embedding. For the test set (N = 13,577), the average difference score was 2.29 amino acid mistakes after reconstruction (standard deviation = 1.54) out of 1,273 amino acid positions.

**Figure 1:**
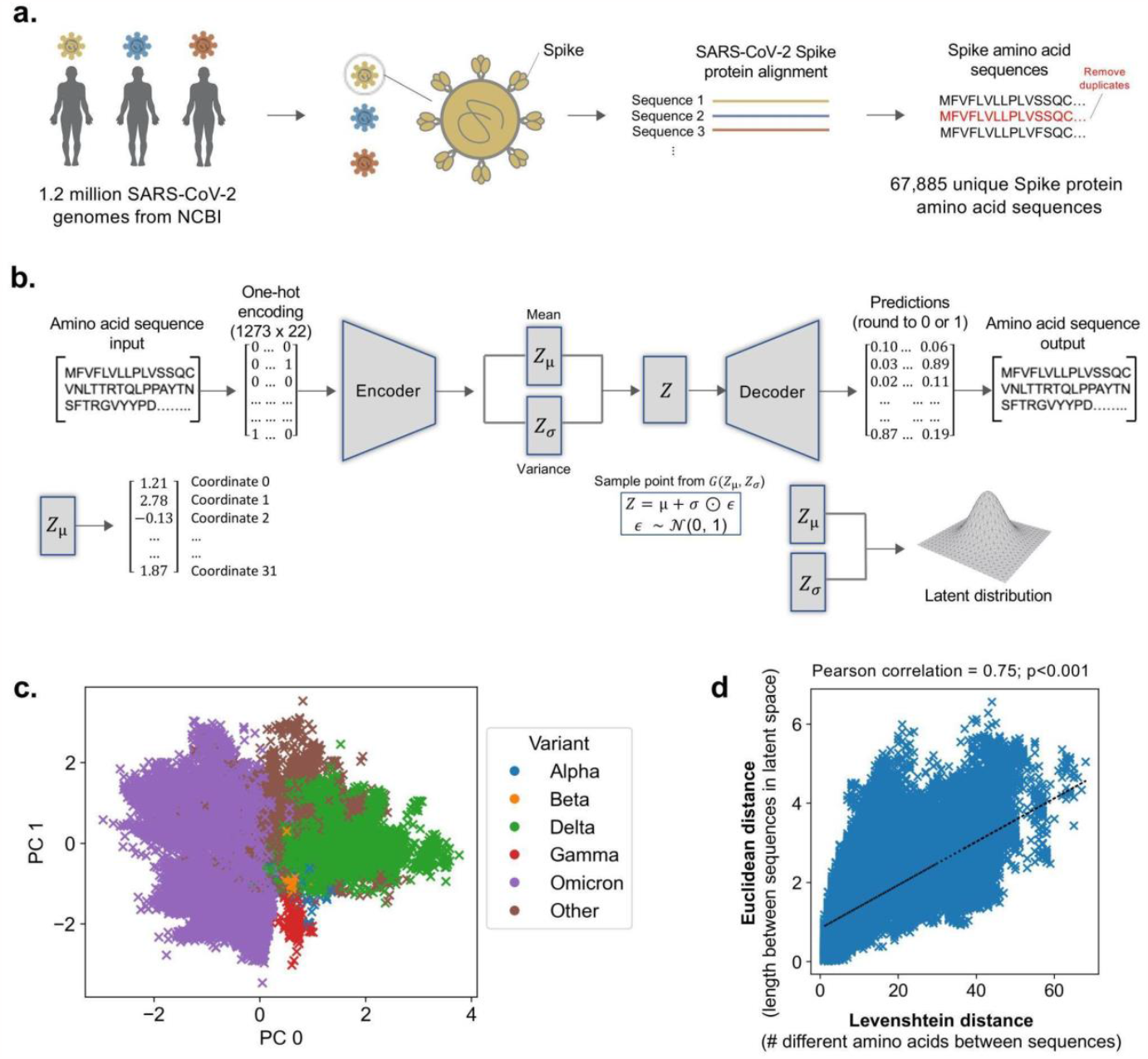
Schematic overview of VAE training and architecture. (**a**) SARS-CoV-2 genomes from the NCBI Virus Database are collected and translated into spike protein amino acid sequences. (**b**) Illustration of VAE architecture. The spike protein amino acid sequences are used to train the autoencoder, where the encoder compresses the sequences into 32 latent space vectors and the decoder reconstructs the encoded sequences back to their original input. (**c**) Scatter plot of the first two principal components of the latent space vectors, colored by SARS-CoV-2 variant. (**d**) Pearson correlation of Levenshtein distances of each sequence pair and Euclidean distances of the corresponding latent vectors, with least-squares line of best fit shown.

To validate the VAE’s accuracy in capturing similarities and differences between spike protein sequences, Levenshtein distances between sequence pairs were compared with Euclidean distances between encoded versions of sequence pairs using Pearson’s correlation (**Figure 1d**). The number of different amino acids between two sequences is represented by Levenshtein distance, while the length of the line segment between two sequences in latent space is represented by Euclidean distance. The two variables were found to be correlated (*ρ* = 0.75, p<0.001), demonstrating the VAE’s ability to accurately capture relationships between sequences in latent space.

### Uncertainty-based neural network prediction of fold changes in neutralization activity against SARS-CoV-2 variants

NCATS OpenData Portal’s curated dataset of 7,069 results from *in vitro* assays was used to train a neural network model to predict fold changes in the neutralization activity of therapeutics and vaccines against SARS-CoV-2 variants (**Figure 2a**). Fold changes are relative to the neutralization activity against the wild-type ancestral Wuhan-Hu-1 strain and were log10 transformed to ensure normality of the ratios. Spike protein sequences of viral isolates subjected to the assays were encoded into 32 latent dimensions using the VAE model, and the 40 therapeutics and vaccines tested against viral isolates were one-hot encoded (**Figure 2b**). The training set comprised data collected between January 9, 2021, and October 31, 2022 (N = 6,089), and the test set comprised data collected between November 1, 2022, and June 22, 2023 (N = 980). We integrated a custom loss function into the model, based on Bayesian inference, to provide reliable estimates of prediction uncertainty (**Figure 2c**). This loss function enabled the estimation of variance associated with each prediction, providing a quantifiable metric that could be used to construct prediction intervals.

**Figure 2:**
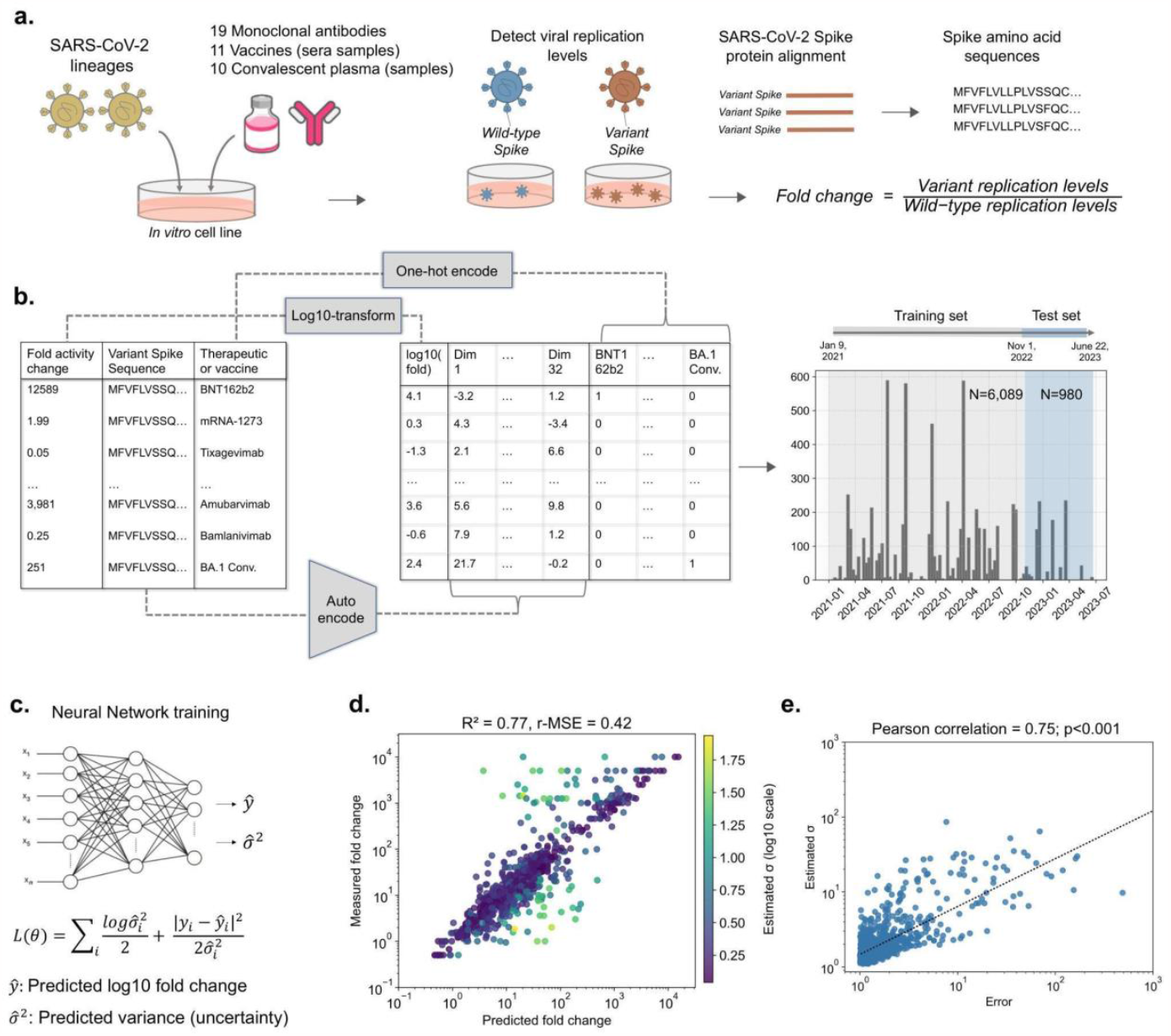
Illustration of neural network training and evaluation for the prediction of SARS-CoV-2 fold change in neutralization activity. (**a**) SARS-CoV-2 isolates were subjected to therapeutic and vaccine *in vitro* assays, and resultant neutralization activity fold change ratios between the wild-type and variants were compiled and log10 transformed. (**b**) Variant spike protein sequences were VAE encoded and corresponding therapeutics and vaccines were one-hot encoded. Data collected from January 9, 2021, to October 31, 2022, was used as training data (N = 6,089), and data collected from November 1, 2022, to June 22, 2023, was used as test data (N = 980). (**c**) A neural network model was trained to predict log10 fold change in neutralization activity and estimate the uncertainty (variance) in each prediction. (**d**) Comparison between model predictions and actual measurements for the test set, highlighting yellow points for higher prediction uncertainty. (**e**) Correlation between prediction error and estimated uncertainty for the test set, with least-squares line of best fit shown.

### Performance evaluation of the uncertainty-based neural network model

Following training, the model demonstrated satisfactory performance on the test set, with an R^2^ of 0.77 and a r-MSE of 0.42 (**Figure 2d**). To ensure that variance estimates serve as a dependable indicator of prediction uncertainty, we compared the absolute prediction error with the estimated variance for the test set using Pearson’s correlation (**Figure 2e**). We observed a statistically significant correlation between the two variables (*ρ* = 0.75, p<0.001), implying that higher prediction uncertainty is associated with a greater likelihood of prediction inaccuracy.

### Assessment of the variability in prediction accuracy

We further evaluated the model’s prediction accuracy across the different COVID-19 therapeutic agents and SARS-CoV-2 lineages. **Figure 3** demonstrates that prediction error varies among the therapeutic agents, with higher mean absolute error (MAE) observed for specific monoclonal antibodies such as Cilgavimab (MAE = 0.62), Bebtelovimab (MAE = 0.55), and S2E12 (MAE = 0.52) (**Figure 3a**). Conversely, prediction error was lower for COVID-19 vaccines and convalescent plasma therapies (**Figure 3b, Figure 3c**). Increased prediction error was also associated with select SARS-CoV-2 lineages, such as CJ.1.1 (MAE = 0.30), BA.2 (MAE = 0.29), and BA.4/5 (MAE = 0.23) (**Figure 3d**). Importantly, it should be noted that estimated uncertainty values for therapeutic agents and viral lineages with larger prediction error exhibited a significant correlation with prediction error, indicating the model’s capacity to provide robust uncertainty estimates when prediction inaccuracy is likely.

**Figure 3:**
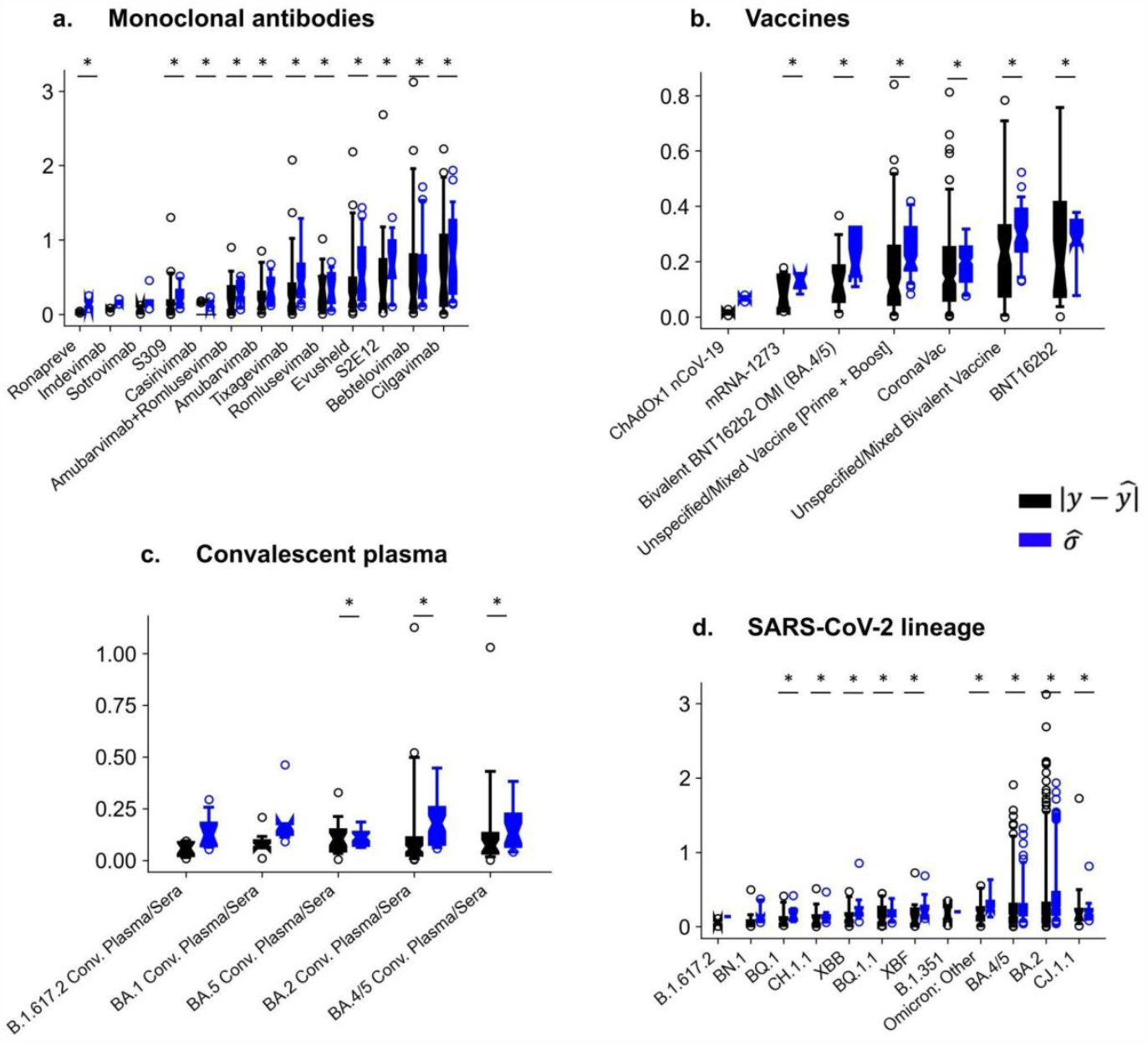
Assessment of neural network model prediction error and uncertainty for COVID-19 therapeutic agents and SARS-CoV-2 lineages in the test set. The distribution of absolute prediction error was assessed for each (**a**) monoclonal antibody, (**b**) vaccine, and (**c**) convalescent plasma sample, as well as for each (**d**) SARS-CoV-2 lineage within the test set, with progressive ordering (left to right) from lowest to highest mean absolute error. Absolute error was compared with the model’s predicted uncertainties 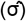 using Spearman’s correlation, with corresponding p-values of less than 0.05 considered statistically significant and annotated within the figures.

### Temporal generalization and performance

We further conducted a temporal analysis to assess the extent to which the model can generalize predictions for prospective variants. Specifically, we evaluated the model’s predictive performance for test sequences assayed within 0-4 months and 4-8 months after the end of the training period of October 31, 2022 (**Figure 4**). The model displayed exceptional predictive capability for sequences assayed 0-4 months after the training period (N = 696), achieving an R^2^ of 0.84 and a r-MSE of 0.34. Despite a decrease in performance, the model still exhibited good predictive capability for sequences assayed 4-8 months after the training period (N = 284), with an R^2^ of 0.64 and a r-MSE of 0.58.

**Figure 4:**
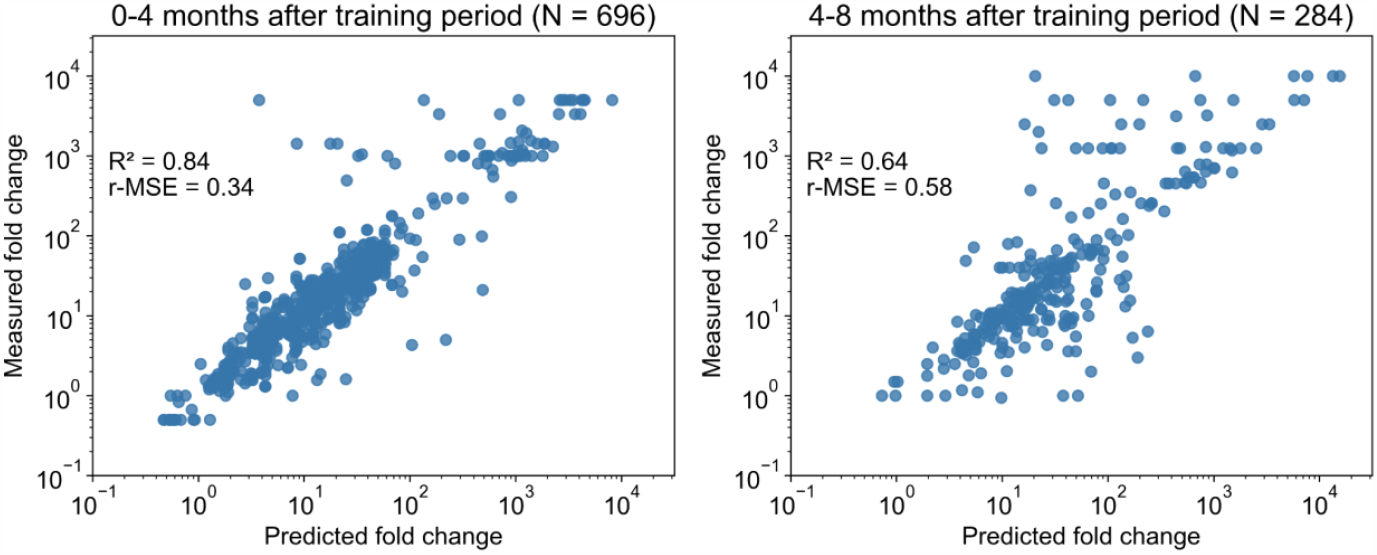
Illustration of the neural network model temporal analysis. Comparison between model predictions and actual measurements for test sequences collected 0-4 months (left column) and 4-8 months (right column) after the end of the training period. R^2^ and r-MSE scores are annotated in each figure.

### Predicted effects for emerging SARS-CoV-2 lineages

As a proof of concept, we present in **Figure 5** the predicted fold changes in neutralizing activity of vaccines and monoclonal antibodies against newly designated SARS-CoV-2 lineages: EG.5, FL.1.5.1, XBB.1.16, and BA.2.86. The predicted effects for parent lineages BA.2 and XBB are also shown as a point of reference.

**Figure 5:**
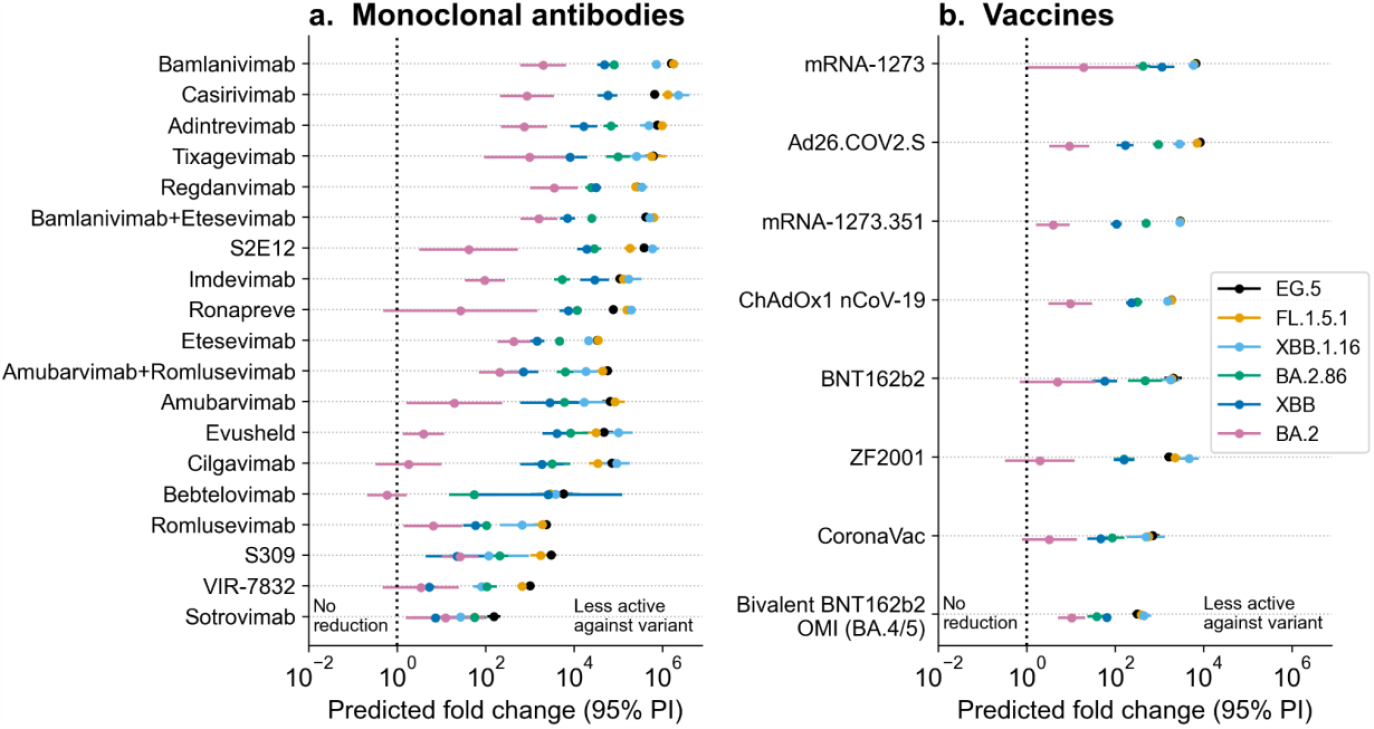
Predicted *in vitro* susceptibility of COVID-19 therapeutic agents to new Omicron sub-lineages. The presented graphs showcase the estimated fold changes in neutralization activity of (**a**) monoclonal antibodies and (**b**) vaccines against the newly emerged SARS-CoV-2 lineages: EG.5, FL.1.5.1, XBB.1.16, and BA.2.86. Parent lineages XBB and BA.2 are also depicted in the figures. Error bars represent the 95% prediction intervals derived from the variance estimates, and the color-coding of the data points and error bars correspond to the lineage to which the prediction was made for.

For this analysis, we trained the VAE and neural network on all available data as of August 18, 2023, to demonstrate the usefulness of our model for surveying new variants in a real-world scenario. The model predicted that both newer XBB descendants (EG.5, FL.1.5.1 and XBB 1.16) and BA.2.86 induce lower neutralizing activity for existing COVID-19 vaccines and monoclonal antibodies, compared to their parental lineages (XBB and BA.2, respectively). However, BA.2.86 is predicted to exhibit a smaller reduction in neutralizing activity to all vaccines and most monoclonal antibodies, compared to newer XBB descendants.

In particular, although bivalent BNT162b2 (OMI BA.4/5) and Sotrovimab are predicted to retain some level of activity against these viral lineages, the predicted fold reduction for bivalent BNT162b2 is 315-fold against EG.5 (95% PI: [247, 401]), 384-fold against FL.1.5.1 (95% PI: [300, 491]), 443-fold against XBB.1.16 (95% PI: [296, 662]), and 38-fold against BA.2.86 (95% PI: [24, 62]). For Sotrovimab, the predicted fold reduction is 155-fold against EG.5 (95% PI: [111, 218]), 56-fold against FL.1.5.1 (95% PI: [30, 106]), 27-fold against XBB.1.16 (95% PI: [8, 97]), and 57-fold against BA.2.86 (95% PI: [30, 106]).

We then conducted a comprehensive analysis of the acquired spike mutations, focusing on identifying those predicted to have the greatest impact on activity of select monoclonal antibodies and vaccines (**Figure S1, S2**). To accomplish this, we first calculated the distinct core mutations distinguishing each new lineage (EG.5, FL.1.5.1, XBB.1.16, and BA.2.86) from their parental lineages. Subsequently, we systematically introduced each mutation, one at a time, into the core spike protein sequence of parent lineage (XBB) to understand the predicted partial impact of each mutation. Notably, the spike mutation F456L, which is present in EG.5 and FL.1.5.1 sequences, is predicted to have the most significant effect on the neutralization activity of therapeutic agents shown in **Figure S1**. Moreover, the largest predicted partial effect of this mutation, relative to XBB parent, is seen for therapeutic agents S309, VIR-7832, and ZF2001. We observed that the newly acquired mutations within the BA.2.86 lineage were generally well-tolerated and did not result in substantial changes in the therapeutic effectiveness (**Figure S2**).

## Discussion

In this study, we outline a novel deep learning approach to monitoring the impact of SARS-CoV-2 variants on the neutralizing activity of COVID-19 antibody therapeutics and vaccines. First, we developed a variational autoencoder (VAE) capable of encoding spike protein sequences into a latent space while preserving the integrity of spike protein information. This resulted in an average reconstruction loss of 2.29 amino acids per 1,273 positions on the spike protein. Subsequently, we trained a neural network model to predict fold changes in the neutralization activity of 40 different COVID-19 antibody therapies and vaccines against spike protein sequence variants, relative to their activity against the ancestral strain (Wuhan-Hu-1). To assess the model’s generalizability to predict prospective sequences, we evaluated its performance on sequences tested eight months after the training data cutoff date of October 31, 2022. Our findings indicate that the model can accurately predict the impact of new spike protein mutants on the neutralization activity of therapeutics and vaccines (R^2^ = 0.77), making it a valuable tool for identifying emerging viral variants that are likely to evade current COVID-19 treatments

Our model’s predictions for emerging SARS-CoV-2 lineages align with current research findings. Consistent with recent data,^27–29^ the model predicts that the newer XBB descendants, specifically EG.5, FL.1.5.1, and XBB.1.16, have significantly reduced *in vitro* susceptibility to both vaccines and monoclonal antibodies, rising concerns for evading vaccine-driven immunity and lack of efficacy to clinically relevant monoclonal antibodies. On further analysis of mutations linked to reduced responsiveness, our model identifies the spike mutation F456L, present in EG.5 and FL.1.5.1, as a primary contributor to reduced neutralization by selected COVID-19 antibodies and vaccines. This observation aligns remarkably with recent studies using pseudovirus neutralization assays,^28,30^ and deep mutational scanning,^31^ which highlighted the mutation’s role in ACE2 receptor binding. The F456L mutation emerged independently in XBB descendant strains such as EG.5, FL.1.5.1, and XBB.1.16.6 and was found in 40% of newly uploaded SARS-CoV-2 sequences by August 2023.^28^ Continuous shifts at this position and its vicinity, highlighted by deep mutational scanning,^31^ suggest the virus is evolving to dodge antibody responses while effectively binding to its target receptors. Remarkably, our deep learning framework successfully discerned the robust immune escape properties of this mutation, despite not seeing it within training sequences for the uncertainty-based neural network, although the mutation was encountered in the VAE training sequences.

On August 17, 2023, the World Health Organization (WHO) designated BA.2.86, a highly mutated subvariant originating from BA.2, as a variant under monitoring due to potential immune evasion risks. However, recent studies employing live and pseudovirus neutralization assays indicate that BA.2.86 elicits a higher neutralizing antibody response to mRNA bivalent boosters than newer XBB descendants,^27,29,32^ though slightly lower than its parental lineage (BA.2). Despite over 30 amino acid mutations in its spike region compared to the parental lineage (BA.2),^32^ many of which are novel to both the VAE and neural network, our model accurately predicted BA.2.86’s behavior, underscoring its performance in emerging lineages. Additionally, consistent with recent studies,^27,33,34^ our model predicted that monoclonal antibodies including Sotrovimab, Cilgavimab, and Bebtelovimab that previously retained some activity against parental BA.2 variants have reduced neutralization activity against BA.2.86. Regarding mutations linked to reduced effectiveness, our findings align with a recent deep mutational scanning study,^31^ which suggests that BA.2.86 mutations are generally well-tolerated and individual mutations do not significantly impact neutralization activities.

Our study showcases the proficiency of deep learning to detect patterns within viral sequence and assay-based data, allowing for the prediction of antibody activity against uncharacterized viral variants. Moreover, we demonstrate the use of Bayesian modeling to quantify prediction uncertainty, enabling the identification of viral sequences and treatments for which the model can confidently make predictions, as well as those with limited assurance where assays would be most informative. The scalability of our approach to monitor therapeutic and vaccine neutralizing efficacy against a wide range of viral strains is particularly beneficial. This can streamline decision-making for the development of immune-based therapies and vaccines, as well as enhance the surveillance of viral strains that may evade current treatments. It can also assist in identifying antibodies that may have regained their effectiveness against emerging variants. As more viral sequence and laboratory-based data continue to accumulate, this approach could be extended to other viral agents, such as influenza and HIV, where antigenic evolution is a critical factor in the development of effective treatments and vaccines. It is also important to highlight that the versatility of this approach extends beyond surface proteins or other specific viral targets, encompassing a broader spectrum of proteins constituting the entirety of the viral genome. In conclusion, this deep learning approach holds promise for addressing the challenges posed by SARS-CoV-2 variants and combating viral evolution in general.

There are several limitations to this study that should be considered. Firstly, the *in vitro* assay data used for model development is subject to experimental bias. Many of the therapeutic agents used to train and evaluate the model are disproportionately represented in the data (**Table 1**). In addition, the number of sequences belonging to specific SARS-CoV-2 lineages within the data set is unevenly spread (**Table 2**). Consequently, these biases may result in reduced model accuracy for select therapeutic agents and lineages, as exemplified in **Figure 3**. It is worth noting that the model does output uncertainty values, which improves prediction reliability and helps address data imbalance issues. Secondly, the performance of our model depends on the accuracy and consistency of previously conducted *in vitro* neutralization assays. While we used fold change ratios to standardize results across different assays, various factors, such as selected cell lines and alterations in viral protein processing, can influence assay readouts and result in inconsistent findings.^35^ Thirdly, it is critical to recognize that the model’s predictions regarding relative changes in *in vitro* neutralizing activity for therapeutics and vaccines do not directly translate to actual treatment effectiveness. *In vivo* studies are indispensable for more precise assessment in this regard, as they encompass a broader array of physiological factors. Lastly, although our model demonstrated satisfactory performance on sequences assayed shortly after the training period, its performance declined for sequences assayed further afterward (**Figure 4**). This suggests that our model is most suited for making predictions for sequences emerging shortly after model training. It is important to note that future endeavors would involve iteratively training the model on all new available data to ensure the most reliable predictions for novel variants.

## Data sources and supplementary material

Sequences used to train the VAE were downloaded from the NCBI Virus Database (https://www.ncbi.nlm.nih.gov/labs/virus/). NCATS, in collaboration with ACTIV TRACE and industry partners, has compiled a dataset of *in vitro* therapeutic activity against SARS-CoV-2 variants from a prioritized set of publications (both preprints and peer-reviewed articles). All variant activity data is made freely available through direct download (https://opendata.ncats.nih.gov/variant/activity). This data was collected on August 18, 2023, and was used to train the neural network model. The autoencoder and neural network models were built in Python (3.10.6). Data was prepared and visualized using NumPy (1.22.4), pandas (1.4.4), and matplotlib (3.5.3). The models were implemented in Keras (version 2.9.0) using a TensorFlow backend (version 2.9.1). The Bayesian Optimization library in python (https://github.com/fmfn/BayesianOptimization) was used to perform a search of the optimal hyperparameters of the neural network model.

## Acknowledgments

The NCATS OpenData Portal program, along with BJP, KW, and KRB’s contributions have been supported by the Intramural program of the National Center for Advancing Translational Sciences, National Institutes of Health and through the Accelerating COVID-19 Therapeutic Interventions and Vaccines (ACTIV) Preclinical Working Group and Tracking Resistance and Coronavirus Evolution (TRACE) initiative with support from the Foundation for the National Institutes of Health (FNIH).

