## Supplementary Material for "A deep learning framework for predicting the neutralizing activity of COVID-19 therapeutics and vaccines against evolving SARS-CoV-2 variants"

### Supplementary Figures:

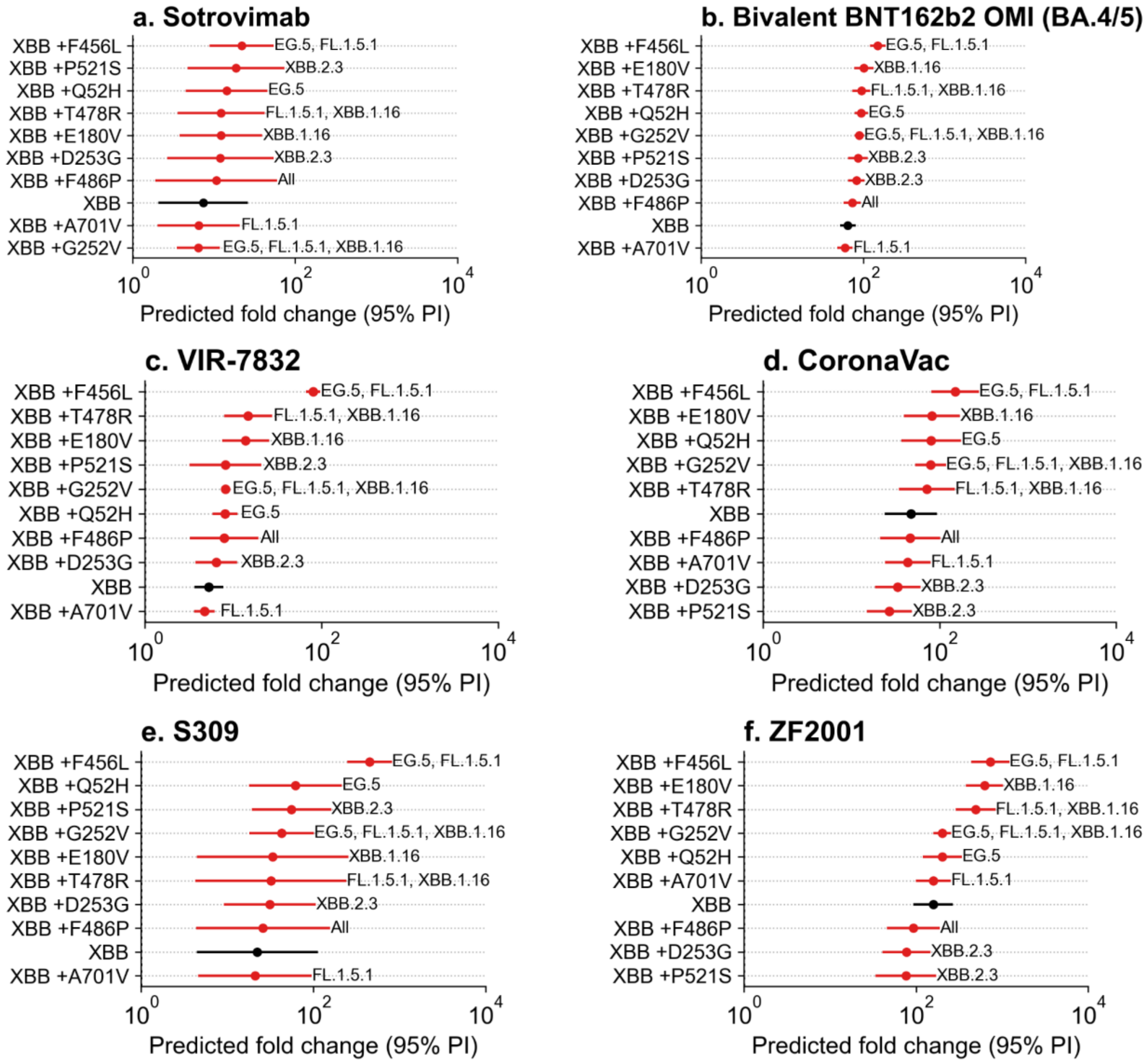


**Figure S1: Impact of specific spike mutations of XBB descendants on predicted neutralization activity changes for selected vaccines and monoclonal antibodies.** The graph illustrates the effect of individual spike mutations within lineages EG.5, FL.1.5.1, and XBB.1.16 on neutralization activity changes. Each unique spike mutation, differentiating from the parent XBB lineage, is systematically introduced to calculate precise fold changes in neutralizing activity. X-axis displays predicted fold change estimates, while the y-axis represents the mutation added to parent XBB lineage. Error bars, illustrating 95% prediction intervals derived from variance estimates.

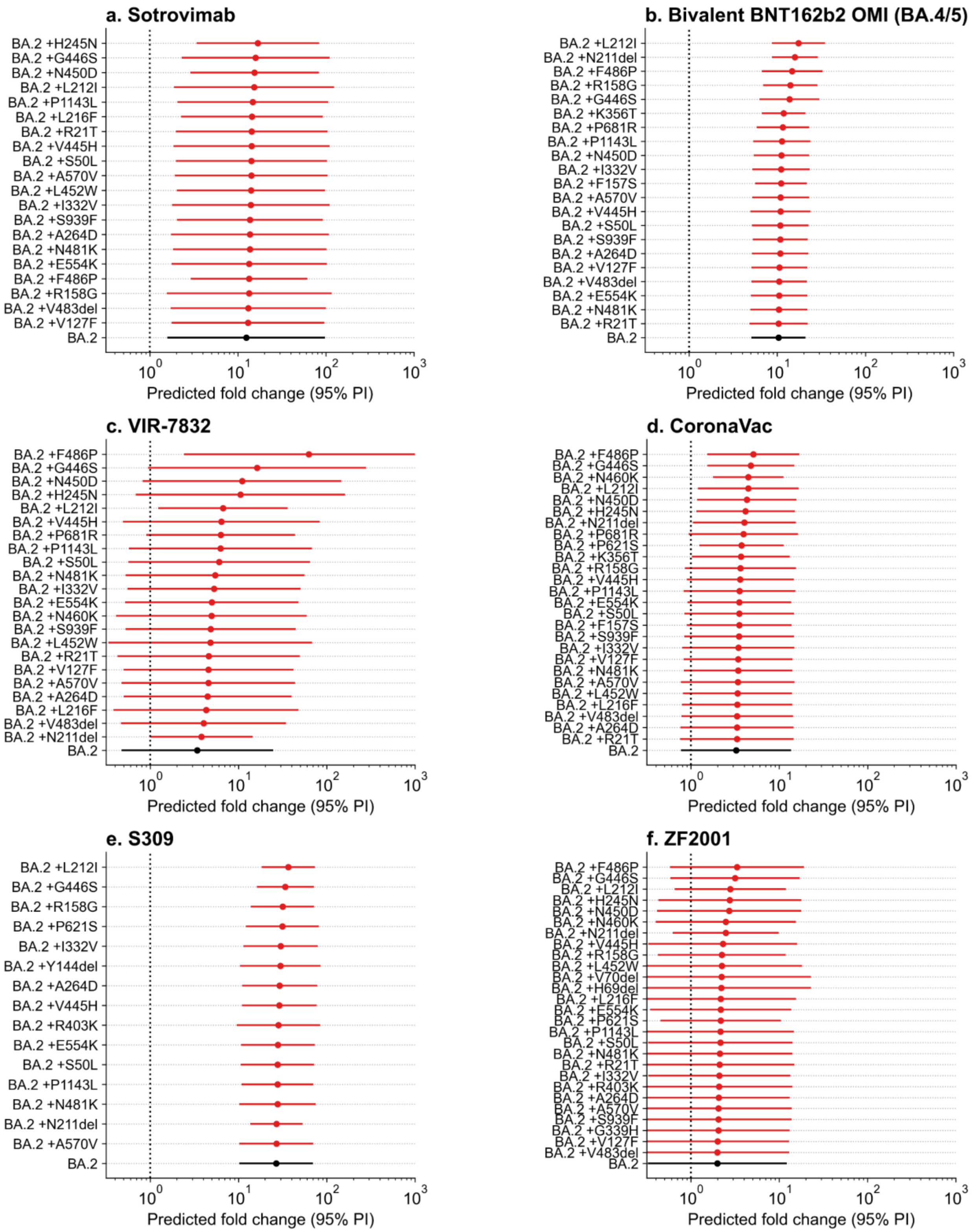


**Figure S2: Impact of specific spike mutations of BA.2.86 lineage on predicted reduction in neutralization activity for selected vaccines and monoclonal antibodies.**
